# Autologous skin cell suspensions established by the VeritaCell method accelerate healing and suppress scarring-associated cutaneous thickening in a rat wound model *in vivo*

**DOI:** 10.64898/2026.03.30.715294

**Authors:** Michael Peake, Olafs Volrāts, Vladimirs Pilipenko, Jolanta Upīte, Arseniy Sergeyev, Baiba Jansone, Nikolaos T. Georgopoulos

**Author notes:** Correspondence: Dr Nik Georgopoulos, E-mail address, Postal address: Biomolecular Sciences Research Centre (BMRC), School of Biosciences and Chemistry, Sheffield Hallam University, City Campus, Howard Street, Sheffield S1 1WB, UK. Joint senior authors (Baiba Jansone, Nikolaos T. Georgopoulos).

## Abstract

Autologous cell suspension (ACS)-based therapies are an established strategy to enhance wound repair, yet limitations in preparation workflows and donor skin requirements remain barriers to wider clinical implementation. We have previously developed VeritaCell, a rapid enzymatic disaggregation-based approach that generates highly viable skin cell populations, including epidermal stem cell-enriched fractions, and demonstrated their pro-regenerative biological properties *in vitro*.

Here, we have evaluated the *in vivo* efficacy of VeritaCell-derived ACS using a rat full-thickness excisional wound model. ACS preparations were applied at donor-to-wound area ratios of 1:1, 1:10, and 1:20, and wound progression was monitored through longitudinal image-based quantification alongside histological assessment of tissue architecture.

ACS-treated wounds exhibited enhanced early wound closure dynamics, with significant within-group improvements evident by Day 6. Histological analysis demonstrated improved neo-epithelial organisation and reduced epidermal thickening in the 1:10 and 1:20 groups, with the 1:10 condition showing tissue architecture most closely resembling unwounded skin. Notably, beneficial effects were observed even at low estimated cell numbers, suggesting that cell viability and biological activity may be key determinants of therapeutic efficacy.

Collectively, these findings provide *in vivo* validation of VeritaCell-derived ACS and support the use of biologically informed donor-to-wound coverage ratios. This approach may enable effective wound repair while minimising donor skin requirements, with potential relevance for the treatment of extensive injuries such as burns.

## Introduction

Breaching of the human skin barrier by insults such as burns, ulcers and physical impact causes damage which normally triggers a complex wound healing response that is driven by the spatially and temporally coordinated action of various cell types. However, wounds that are either too large (burns) or non-healing (diabetic ulcers) recover more slowly and often may be associated with microbial infection which curtails the effectiveness of clinical wound management, thus resulting in enormous financial burden for healthcare providers. For the National Health Service (NHS) in the UK alone, the cost of wound management is now estimated to be £8.3 billion (1).

For the treatment of large wounds (such as burns), in most cases split-thickness skin grafting (STSG) is utilised, whereby an autologous skin graft is taken from the patient and placed onto the wound site area. Yet, this method can demonstrate inefficiency in healing, poor aesthetics and lengthy hospital stays due to large amounts of donor site skin used (2). Thus, there is an urgent clinical need for producing novel, alternative approaches to achieve rapid and efficient wound recovery and improvement in patient wellbeing. A number of alternative approaches have been developed, which include: a) use of engineered tissues as skin ‘substitutes’ that primarily function as barriers for prevention of fluid loss and microbial contamination (3); b) isolation and extensive ex vivo expansion of skin cells (keratinocytes and fibroblasts) for subsequent use in various procedures directly, such as allogeneic cell therapy (4), or by their incorporation into scaffold materials (5), or in three-dimensional bio-printed constructs for creation of physiologically functional skin structures (6). Nevertheless, STSG still represents the current method-of-choice in wound care, because, despite their promise, the aforementioned novel strategies have been unable to contemporaneously offer improvements in cost-effectiveness, ease of practical use, and efficacy compared to STSG.

A promising alternative strategy that can overcome such obstacles is a cell-based, “point-of-care” therapy which involves application of autologous skin cells. This involves enzyme-based isolation of cells from a patient’s own skin *in situ*, which can then be immediately applied to the wound area (7). An advantage of such autologous cell suspension (ACS) is the requirement of a much smaller amount of donor skin. Unlike STSG, which requires donor to wound site ratios of 1:1 or 1:2, there is evidence that ACS application may instead require remarkably less donor skin (8). However, although the concept of ACS-based therapy is promising, it has shown inadequate clinical success, and the costs of using currently available devices for ACS isolation remain extremely prohibitive.

To overcome the limitations of the currently available methods and devices for preparation of ACS isolates, we have recently established VeritaCell for rapid, enzymatic disaggregation of human skin cells. Using this methodology, which requires no specialist instrumentation and is performed at room temperature, we have reported efficient recovery of keratinocytes, fibroblasts and melanocytes. VeritaCell ACS-isolates contained Epidermal Stem cells (EpSCs) and exhibited a wound healing-enhancing secretome which accelerated keratinocyte proliferation and markedly reduced cytodifferentiation of myofibroblasts, the mediators of scarring (9). Thus, our previous *in vitro* studies provided several lines of evidence that the established ACS isolates possess biological properties that may enhance the speed and the ‘quality’ of wound healing.

To provide functional evidence that when such ACS suspensions are applied to wounded skin *in vivo* can enhance neo-epithelialisation and improve the quality of the wound healing process overall, here we tested our methodology using the rat excisional wound model. To assess the effect of the suspensions on such acute wounds, we employed a series of donor-to-wound size ratios to unravel possible differences in efficacy, determined the rates of wound closure (neo-epithelialisation) following cell application, as well as examining the quality of wound recovery by assessing the organisation and uniformity of the neo-epithelium and observing for scarring-associated thickening.

## Materials and methods

### Animals and ethics

We recruited 32 male *Wistar* rats weighing 220-240g, obtained from Charles River Laboratories (Sulzfeld, Germany). The animals were housed in environmentally enriched cages (4 rats per cage), located in individually ventilated stainless-steel racks (GR900, Tecniplast, Italy). Each cage contained autoclaved aspen wood chips (1031004, LBS-Biotech, UK) in addition to enrichment items including a polycarbonate tunnel (K3325) and aspen blocks (1023007) obtained from LBS-Biotech (UK). Rooms were maintained in accordance with experimental animal welfare regulations, under standard laboratory conditions (temperature 23°±1C; humidity 50-60%, 12-h day/night cycle 7:00 to 19:00; cage environmental enrichment) throughout the study. Rodents received a standard cereal-based pelleted chow diet (19.2% protein, 4.1% fat, 6.1% fibre, and 5.9% ash) (1324, Altromin, Mucedola, Italy) and filtered tap water supplied *ad libitum*. Rats were allowed to habituate to the environment of the animal facility for 9 days (Figure 1) before experimental procedures started.

**Figure 1.**
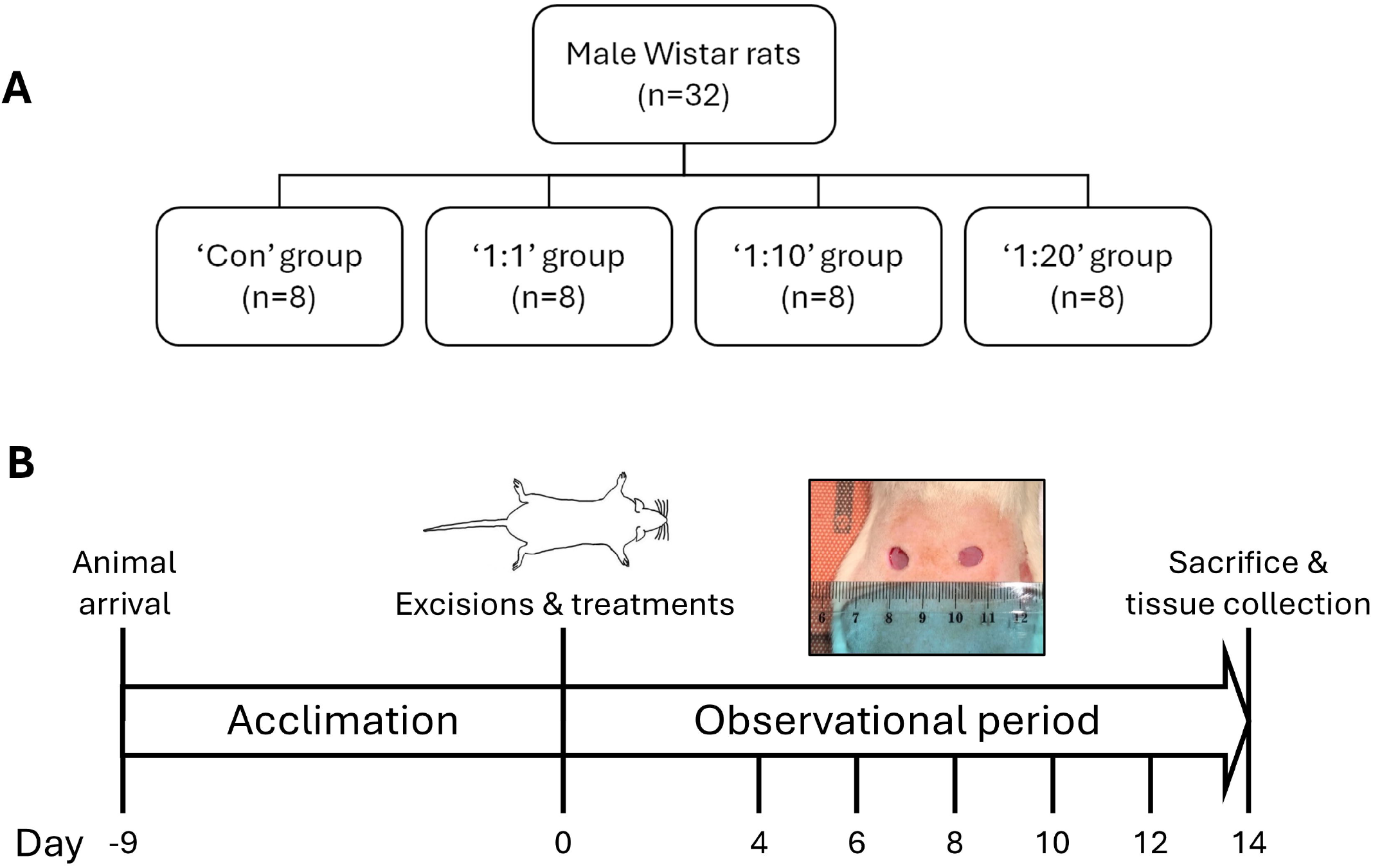
Study design. Schematic representation of the work plan to assess the effect of ACS suspensions isolated using the VeritaCell methodology on acute wound healing *in vivo*. **A)** Control and test animal groups. **B)** Timeline of the *in vivo* experiments.

The animal protocol for this study was approved by the Local Animal Welfare Committee of the University of Latvia and Animal Ethics Committee of the Food and Veterinary Service, Riga, Latvia (Permit number: 135/2022) and all experimental procedures (e.g. study design, surgical manipulations, postoperative care and humane endpoints) were performed in accordance with the EU Directive 2010/63/EU and local laws and policies on the protection of animals used for scientific purposes. All efforts were made to minimize animal suffering and reduce the total number of animals used for this study.

### Chemicals

Isoflurane was obtained from Vetpharma Animal Health (Spain) and buprenorphine from Le Vet Beheer (Netherlands). Ethanol (70%) was supplied by Kalsnava distillery (Latvia) and Ringer’s solution was procured from Fresenius Kabi (Poland).

### Experimental design

To assess the ability of ACS to enhance wound healing, we tested (alongside Ringer’s solution-treated Controls) three different ACS cell suspension densities (Figure 1). Prior to the establishment of excisional wounds by punch biopsy (section 2.4), a 1-2 cm^2^ STSG was removed using a dermatome from the dorsal skin of each animal and ACS solutions were prepared using the VeritaCell methodology (Figure 2). The density of each sample was defined as the ratio between the size of the donor site *versus* the size of the treatment area. More specifically, we tested ratios 1:1, 1:10 and 1:20 (by appropriate dilution of the original ACS) with 1:20 indicating coverage of a wound area 20x larger than the donor site.

**Figure 2.**
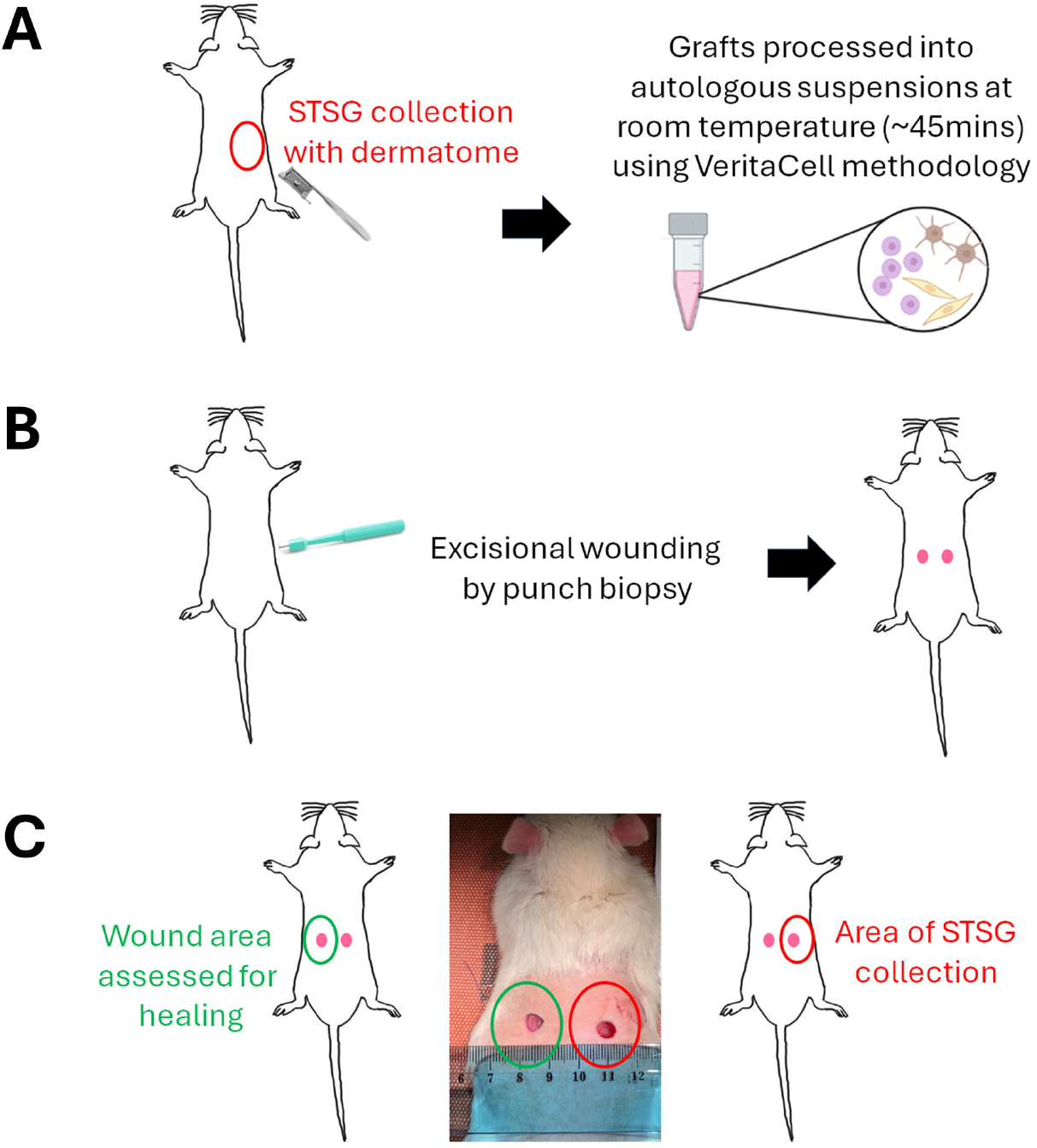
Preparation of ACS isolates for application on wounds. **A)** STSG was excised using a dermatome from the right side of the dorsal area of each animal and skin was processed using the VeritaCell methodology for the isolation of ACS suspensions. **B**) Excisional wounds were established using a punch biopsy tool. **C**) As the punch biopsy method generates two symmetrically positioned wounds on the dorsal area, both wounds were treated with either ACS isolates or control solution. However, the donor STSG collection area (wound indicated in red) was only utilised to perforate the skin, and it was the opposite wound (denoted in green) that was used to assess healing rates.

For this purpose, rats were randomly divided into the following 4 treatment groups (n=8 in each group): control, 1:1, 1:10 and 1:20. The treatments were applied on each wound only on days 0 (day of wound induction). After treatment administration, wounds were covered with dressing, bandage and the patch until day 4 (see below). Wound evolution was monitored by wound imaging on days 0, 4, 6, 8, 10, 12 and 14. The animals’ general health status was observed twice daily during the study period.

### Excisional wound model induction

A day before wounding, the dorsal fur of the animals was shaved with an electric clipper under anaesthesia and disinfected with 70% ethanol. On the day of wounding, rats were weighed and to prevent acute pain buprenorphine (0.05 mg/kg, s.c.) was injected 40 min before the surgery. Anaesthesia was induced via 4.5% isoflurane, maintained with 1.5% to 2% isoflurane in 0.3 L/min O_2_ and 0.7 L/min N_2_O, using a facemask anaesthesia system. The dorsal skin of the rat was stretched along the midline of the spine and two circular full-thickness wounds were created with a disposable 8 mm round skin biopsy punch tool (Kai Medical, Japan) symmetrically along the dorsal midline of rats (10). The formed wounds were rinsed a few times with saline. A total of two wounds were created on each rat back and to avoid cross contamination both wounds were administered with the same treatment. Before the application of the treatment, wounds were washed with saline and then the treatment (Control or VeritaCell-derived cell suspensions) in a volume of 100μL were applied on both wounds (Figure 2) of each animal. Approximately 30 seconds after the application of the treatment, wounds were covered with one layer of a 4 cm x 4 cm piece of nonadherent, low-absorbent, small pore dressing from Telfa™ Clear Wound Dressing (Covidien, Minneapolis, MN) to protect the wound area from potential infection and was followed by an elastic self-adhesive bandage Nowopress (KOB GmbH, Germany). Finally, a porous, elastic patch Mefix (Mölnlycke Health Care, Sweden) was wrapped around the trunk of the animal to provide a secure hold. After dressing, rats were injected subcutaneously in the back with 2 mL Ringer’s solution. After full recovery from anaesthesia, each animal was housed in an individual cage until the end of the experiment (14 days) to avoid communal licking of wounds. The dressing, bandage and patch were removed on day 4 of the study and rats were left undressed to the open environment of their home cages.

### Macroscopic determination of wound closure rates

To observe the recovery of the wound areas, wounds were photographed using a digital camera on Days 0, 4, 6, 8, 10, 12 and 14. Images were acquired using an iPhone SE (3rd generation) equipped with a 12 MP wide-angle rear camera (ƒ/1.8 aperture, 5x digital zoom, and Smart HDR 4). All images were captured under consistent lighting conditions using the same resolution (4032 × 3024 pixels) and standardized camera settings to ensure uniformity across all time points and treatment groups. Following wound image collection, the size of each wound area was measured using ImageJ software (National Institutes of Health, Bethesda, MD, USA). The wound areas were determined by calculating the ratio of the initial area to the wound area at various time intervals. The wound margins were traced and the % wound closure at each time point was calculated (in pixels) using the following equation and as elsewhere (11): Wound healing rate (%) = ((Wound area on day 0 - Wound area on day(n))/(Wound area on day 0)) × 100%. All image analyses were carried out by a reviewer blinded to the experimental groups.

### Histology

Tissues samples from healing wounds were collected from all animal groups under deep anaesthesia. Each wound tissue sample was excised with a 0.5 cm safety margin of the entire skin around the initial wound area by using scissors, with depth to the muscular fascia. The removed tissue was immediately placed in 10% (v/v) formaldehyde solution and fixed for further histological analysis. Skin samples were embedded in paraffin wax, 4 μm tissue sections prepared, dewaxed and Hematoxylin & Eosin (H&E) staining was performed using a Dako Reagent Management System (DakoRMS) and by automated processing (dehydrating, cover-slipping and slide-drying) on a Dako CoverStainer (Agilent).

### Assessment of mean epidermal layer thickness

Histochemistry microscopy images of wound tissue specimens (above) were analysed to quantify epidermal thickness across treatment and control groups using ImageJ® software (https://imagej.net/). Any images that demonstrated experimental ‘noise’/artifacts, and thus did not permit full epidermal layer length to be analysed, were excluded from the analysis to avoid any bias. Within each available image, evenly spaced lines were drawn across the whole of the epidermis (to determine the distance between the basal boundary of the *stratum basale* and the apical boundary of the stratum corneum) within the image field of view (a minimum of 30-35 lines drawn in total per image). The lengths of these lines were measured and averaged to determine the top-down mean epidermal thickness.

### Statistical analysis

All experimental data are expressed as the means ± standard deviation (S.D.) or standard error of the mean (SEM). Results from quantification of wound samples were averaged for each wound site, the means were then used for analysis using GraphPad Prism version 8.3.0 (GraphPad Software, San Diego, California). Comparisons between multiple groups were made using one-way or two-way ANOVA with Tukey’s post-hoc test. *P*-values□≤□0.05 were considered statistically significant (individual p values within different treatment groups are appropriately indicated in the relevant figure captions).

## Results

### Study design to assess the ability of ACS isolated by the VeritaCell methodology to enhance acute wound healing responses *in vivo*

We have previously demonstrated that our VeritaCell methodology permits isolation of highly viable human epidermal keratinocytes and fibroblasts from small STSG samples, and shown that such ACS isolates contain EpSCs that are known to support epidermal re-epithelisation and enhance the wound healing response. As the method involves enzymatic disaggregation of skin as well as filtration steps for isolation of cell populations (9), we sought to test the safety of the methodology and its ability to influence the process of wound healing *in vivo*.

We employed the rat excisional wound model to test the effect of our ACS on such acute wounds. As a key aim of using ACS from a small donor skin piece is to be able to ‘cover’ a several fold larger wound area, we tested different ‘concentrations’ of such ACS isolates, to mimic different wound coverage ratios. Specifically, using an approximate ∼1-2 cm^2^ donor skin size, we tested whether such isolates could benefit wounds that were 10x (‘1:10’ ratio) or 20x (‘1:20’ ratio) larger in surface area than the donor site. We tested these alongside wounds treated with the equivalent of a full suspension (‘1:1’ ratio) and compared ACS-treated wounds to untreated (‘Control’) wounds (Figure 1).

Of note, as the punch biopsy method generates two symmetrically positioned wounds on the dorsal area, both wounds were treated with either ACS isolate or control solution. However, as schematically represented in Figure 2, the side from which donor STSG was provided to establish cell suspensions was only utilised to perforate the skin, and it was the opposite wound that was used to assess healing (as detailed in the figure caption).

### Treatment of acute wounds with ACS suspensions enhances wound healing as determined by measurement of % wound closure rates

Following establishment of punch biopsy-induced wounds and treatment with the three ACS densities (1:1, 1:10 and 1:20) or control solution (Day 0), wounds were covered with low adherence dressings (to minimise ‘damage’ to the applied cell populations) and were left undisturbed for 4 days. Dressings were then removed, and all wound sites were imaged every 2 days thereafter until Day 14, and representative images are provided in Figure 3. Through visual inspection, it was observed that the wound tissue was in good condition in both the control and treated groups, and no signs of inflammation or wound deterioration were evident during the healing period. Yet, interestingly, we observed the clear presence of tissue exudate and a small amount of bleeding on Day 4 for Control wounds. By contrast, all wounds treated with ACS exhibited no such exudate and were visibly healthier sooner. By Days 6 and 8, however, no such exudate was observed in the Control group (Figure 3), too.

**Figure 3.**
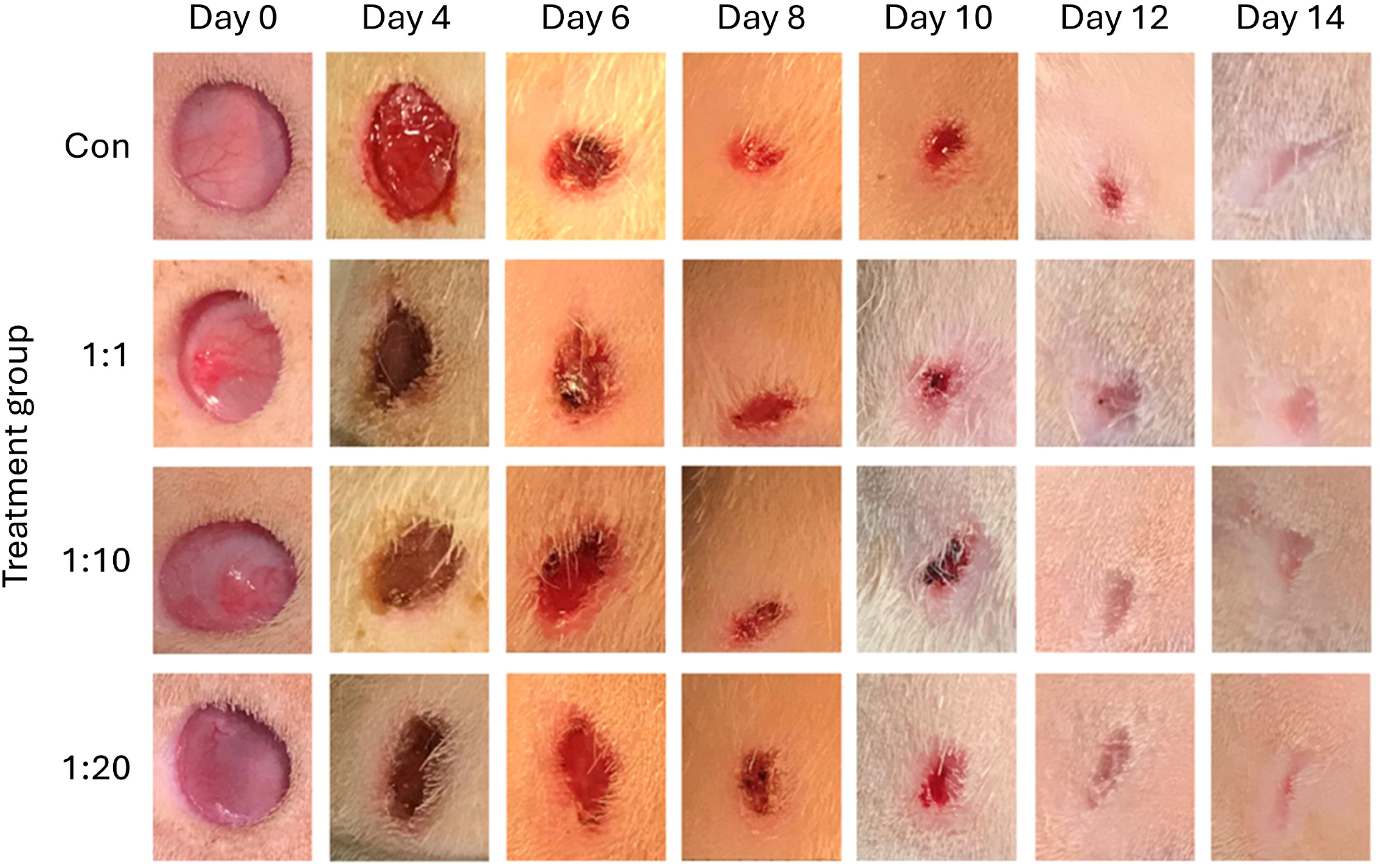
Imaging of wounds at different time points post-treatment. Excisional wounds were imaged on the way of establishment, prior to treatment with ACS suspensions or control solution (Day 0) and application of dressings (as detailed in the Methods). Following a period of 4 days during which wounds were undisturbed, dressings were removed (Day 4) and wounds imaged every 2 days. Representative images of wounds throughout the 14-day assessment period are provided.

Wound area image analysis (carried out as presented in Figure 4A) permitted determination of % wound healing rates, as shown for all groups per day of assessment in Figure 4B. Although our analysis revealed statistically significant changes between days (p<0.0001), wound healing rate values did not significantly differ between groups (p=0.91), indicating that for these acute, rapidly-healing wounds no significant changes in healing rates existed between groups on the days tested. However, further analysis allowed more subtle yet consistent differences to be determined when the data was assessed per treatment group.

**Figure 4.**
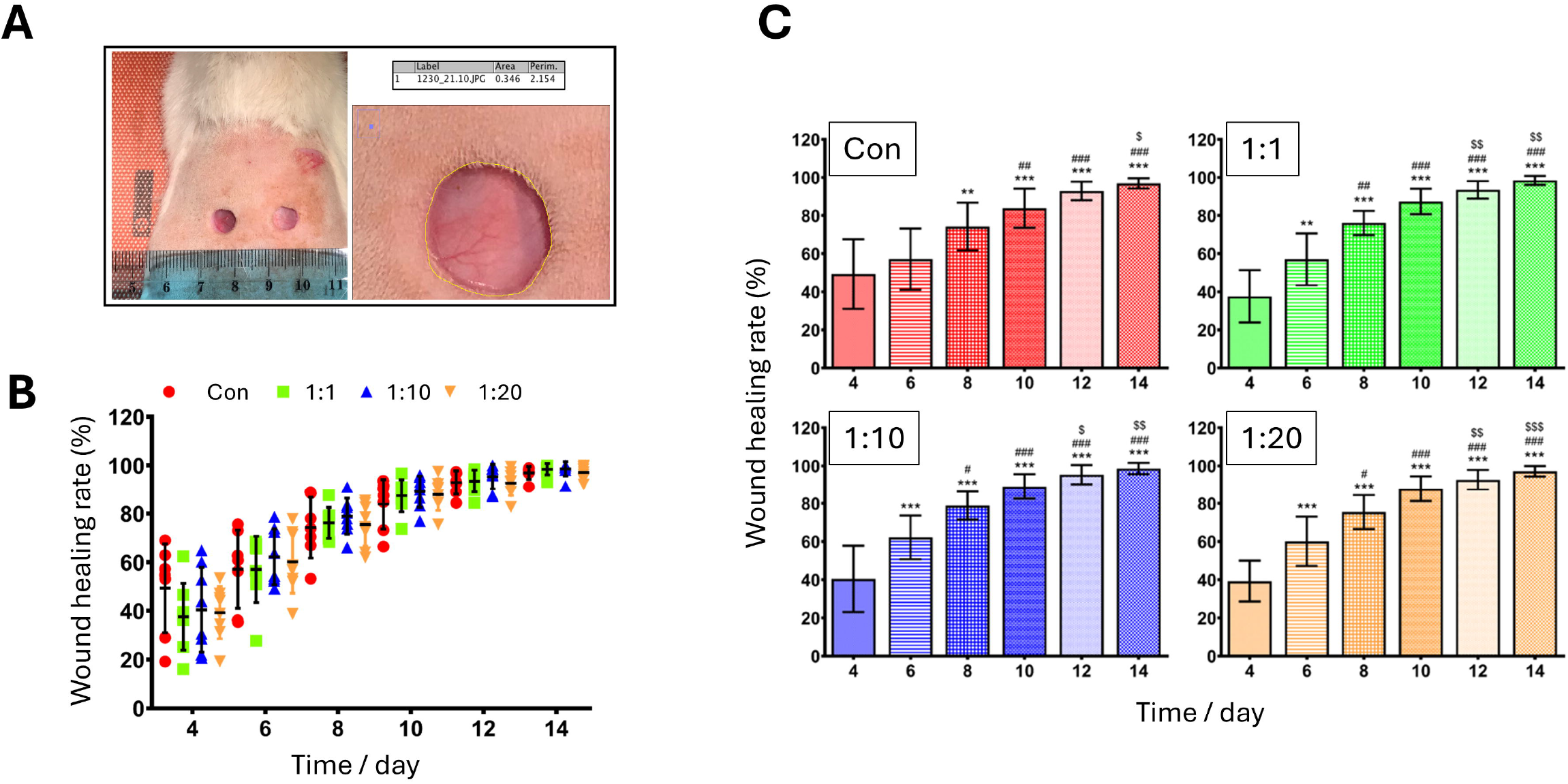
Assessment of wound closure (% healing rates). **A)** Using ImageJ, wound size area at each time point was determined as indicated using the software drawing tool and following ‘calibration’ of image resolution. For each treatment (Control, 1:1, 1:10 and 1:20), the measurement was repeated 3 independent times, and a mean wound ‘Area’ was calculated. **B)** Wound closure rates (expressed as ‘% wound healing rate’) for all groups during the 14-day period were calculated (as detailed in the Methods). Values are shown as mean ± S.D., and all individual values are presented for each group (n=7-8 per group). **C**) Wound healing rates are presented for individual treatment groups for the purposes of denoting the results of statistical analysis. Data represent mean ± S.D. (n=7-8 per group). ^**^p<0.01, ^***^p<0.001 vs. Day 4; ^#^p<0.05, ^##^p<0.01 and ^###^p<0.001 vs. Day 6; ^$^p<0.05, ^$$^p<0.01 and ^$$$^p<0.001 vs. Day 8.

As shown in Figure 4C, when % healing rates were analysed within each individual group, significant changes were observed and, more importantly, these patterns of significance differed between control *versus* treatment groups. More specifically, in the Control group a significant increase in % healing rate was observed on Days 8, 10, 12 and 14 when compared to Day 4 (p<0.0001 for each); notably, however, there was no significant difference for Day 6 when compared to Day 4. By contrast, in all treatment groups, a significant increase in % wound healing rate was already observed on Day 6 when compared to Day 4, as indicated by p<0.01 for the 1:1 group, and even more significantly by p<0.001 for both 1:10 and 1:20 treatment groups (Figure 4C). Furthermore, % healing rates in all treatment groups were significantly higher on Day 8 compared to Day 6 for groups 1:1, 1:10 and 1:20 (p<0.001), in comparison to lower significance in % healing for Day 8 compared to Day 6 for the Control group (p<0.01). Interestingly, particularly notable was that the 1:10 treatment group exhibited higher % wound closure at Day 6 and was the first treated group to exceed 80% wound closure by Day 8 (Figure 4C).

### Histological assessment of neo-epithelialisation of ACS-treated wounds reveals an enhancement of the quality of wound healing and a reduction in scarring-associated cutaneous thickening

Following completion of the wound closure experiments, tissue was collected from the wound sites and analysed by histological examination to assess the quality of the wound recovery. Inspection of the tissue architecture from representative section images shown in Figure 5, revealed striking differences between treatments. Control wounds (that were not treated with cell suspensions) showed a dense and relatively disorganised extracellular matrix in the dermis and extensive thickening of the epidermis, which was clearly highly-variable in its thickness thus indicative of a scarring-associated response. On the other hand, all wounds treated with ACS isolates exhibited better ‘quality’ of neo-epithelialisation.

**Figure 5.**
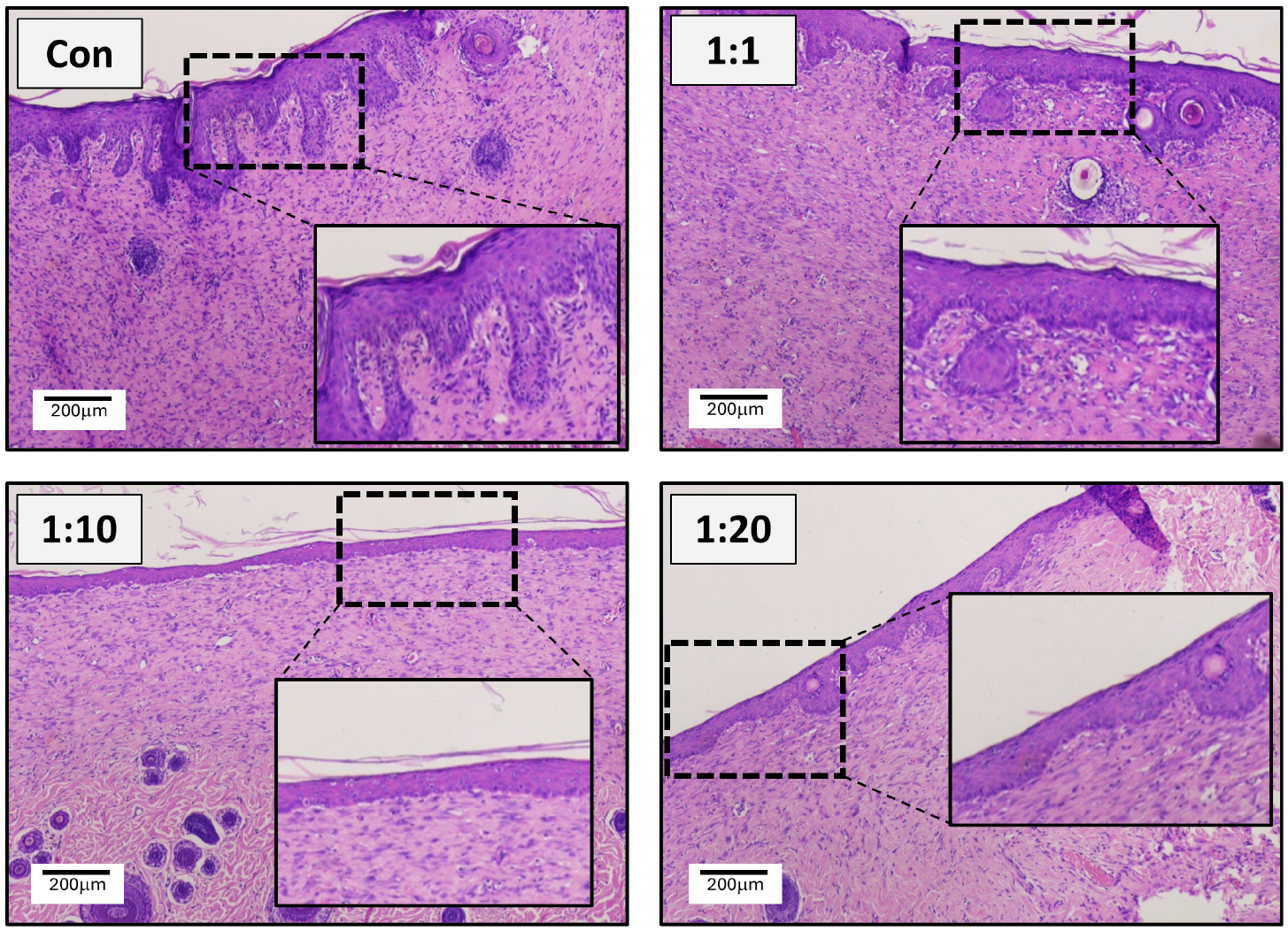
Histological examination of wounds. Following completion of the experiment time-course, on Day 14 all animals were sacrificed and wound tissues were collected for H&E staining. Representative images from all treatments (n=7-8 per group) are shown, and image inserts are included to provide clearer representation of neo-epithelialisation. Scale bar: 200 μm.

Interestingly, despite an improvement in the architecture of both the epidermal and dermal areas, 1:1 treatment was still variable in its thickness. By contrast, both the 1:10 and 1:20 treatments showed a well-organised underlying dermis and exhibited an epidermal layer that was visibly thinner and less variable in its thickness (Figure 5), with the 1:10 group demonstrating particularly reduced thickness and minimal variability (similar to unwounded intact skin). Furthermore, to provide an objective assessment of epidermal layer thickness in the treatment groups, we comprehensively assessed the epidermal layers and determined mean layer thickness (as indicated in Figure 6A). The results shown in Figure 6B, confirmed that all treated wounds showed reduced neo-epithelium thickness, which was evident for the 1:20 group and was particularly striking for the 1:10 group (although the result did not reach statistical significance – adjusted p-value 0.07 using Kruskal-Wallis analysis).

**Figure 6.**
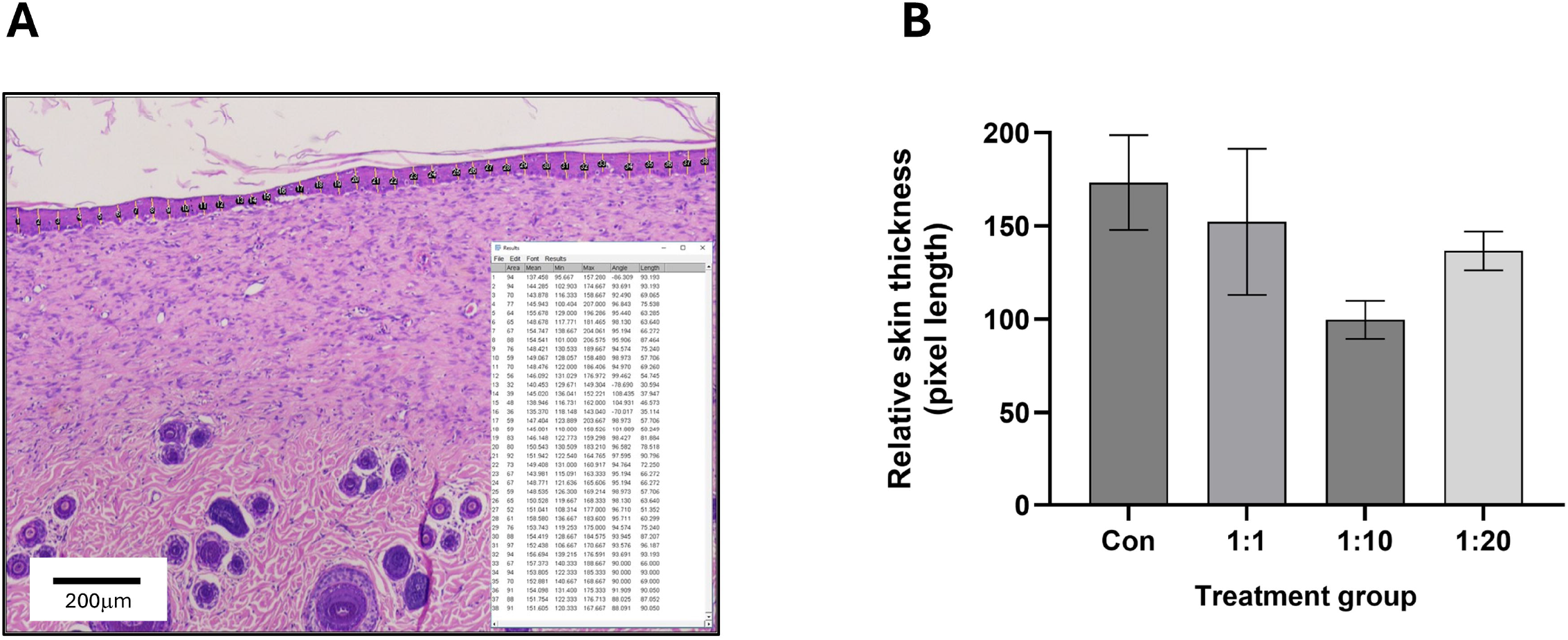
Determination of epidermal layer thickness in wounds. **A)** Microscopy images of wound tissue histochemistry were analysed to quantify epidermal layer thickness across all treatment groups using ImageJ (as detailed in the Methods). As indicated in the representative image, evenly spaced lines were drawn to determine the distance between the basal boundary of the *stratum basale* and the apical boundary of the *stratum corneum* across the whole of the epidermis within the field of view. The lengths of the lines were measured and averaged to determine the top-down mean epidermal thickness. **B)** Relative skin thickness (pixel length based) across treatment and control groups (as in Figure 5) was calculated and data are expressed as mean relative skin thickness ± SEM.

## Discussion

Unlike the normal process of skin re-epithelialisation by the rapid proliferation and migration of keratinocytes, deceleration of neo-epithelialisation is associated with delayed tissue repair, infection and/or chronic wound development. It is thus essential to provide solutions that isolate viable cell populations which can then be applied directly and enhance the process of epidermal regeneration. There is increasing evidence that ACS-based therapies represent a highly-promising approach (7, 12), with clinical success for burns (8, 13) and some promise for chronic wounds (14, 15). ACS-associated cells integrate (“take”) into a wound site (16, 17), which accelerates re-epithelisation and wound healing, reduces infection risk and suppresses discomfort (pain). However, despite the promise of the approach, existing technologies for application of ACS have not achieved the desired clinical success due to several limitations (9, 18), whilst the ability to obtain optimal cell populations often requires electrical and/or laboratory instrumentation (19).

To address these limitations, we have recently established VeritaCell, a novel methodology for rapid, enzymatic disaggregation of human skin cells and establishment of ACS isolates, and we examined the biological properties of these cell populations. We reported extremely efficient recovery of keratinocytes, fibroblasts (and melanocytes) and VeritaCell ACS-isolates contained Epidermal Stem cells (EpSCs) that were demarcated via the CD49-high/CD71-low protein expression profile. The isolated cells exhibited a wound healing-enhancing secretome which both accelerated keratinocyte proliferation and markedly curtailed cytodifferentiation of myofibroblasts, the key mediators of fibrosis and scarring (9). Thus, our *in vitro* studies previously implied that the established ACS isolates could support wound healing and possess biological properties that have the potential to enhance not only the speed but, equally, the ‘quality’ of wound healing. To provide functional evidence that when such ACS suspensions are applied to wounded skin *in vivo* they can, based on the above biological properties, enhance neo-epithelialisation and improve the quality of the wound healing process overall, here we tested our methodology using the rat excisional (punch biopsy-based) wound model.

Amongst a variety of models for cutaneous wound healing assessment (20), excisional wound models in rats are widely used to study the biological processes of skin repair, despite anatomical differences between rodent and human skin. These models involve circular 8–20 mm full-thickness skin wounds (excisions) to investigate phases of healing, including inflammation, granulation tissue formation, re-epithelialization, and remodelling (21). For wound establishment, full-thickness excisions are created on the dorsum using punch biopsy tools and wound evaluation metrics utilised include macroscopic wound area reduction tracking (via callipers or imaging) and subsequent microscopic, histological scoring of epithelialization as well as inflammation, angiogenesis, and collagen deposition (22). Here, we created such wounds in the dorsal area and assessed the effect of VeritaCell ACS isolates at different donor-skin-size *versus* wound-size ratios on wound healing.

Even at the early stages of wound healing, ACS-treated wounds showed reduced exudate levels and visually demonstrated more rapid recovery. Untreated (Control) wounds exhibited gradual closure within a 14-day follow-up period and significant healing (% wound closure) was only observed after Day 8. By contrast, treatment of wounds with ACS suspensions of 3 different concentrations (1:1, 1:10 and 1:20) led to significant % wound closure within the first week (Days 6 & 8). Treatment with 1:10 suspensions showed particularly marked improvement in wound closure which was the most significant throughout the healing stages. Albeit statistically significant, one may argue that these improvements may appear relatively modest. However, it must be noted that we assessed the effect of ACS suspensions in acute and naturally rapidly-healing wounds. Yet, even in such a biological context, by providing high numbers of viable skin cell populations, VeritaCell suspensions still significantly enhanced wound closure during the critical, early-stages of the wound healing response.

Histological assessment of wounds provided even more notable observations. Untreated wounds on Day 14 showed a dense and visibly disorganised extracellular matrix network, and most notably exhibited extensive thickening of the neo-epidermis, which had ‘invaded’ deeply into the underlying dermis and was highly-variable in its thickness, observations confirming marked cutaneous thickening and indicative of a scarring-associated response. Instead, upon treatment with ACS, there was improved and smoother organisation of the underlying matrix in the dermis. For both 1:10 and 1:20 treatments, the epidermal layer that was thinner and far less variable in its thickness, as confirmed by detailed image analysis. Particularly striking were the observations for the 1:10 group, which demonstrated reduced thickness and such minimal variability that it was essentially histologically identical to unwounded skin. Interestingly, although one might have anticipated that the 1:1 ratio would have provided the best response (potentially due to the provision of higher ACS cell density), clearly the additional cell numbers did not necessarily translate to superior enhancement of the wound healing process. To our knowledge, this is the first such observations for ACS-based treatment of wounds. It is tempting to speculate that, these observations may be indicative of the importance of supporting the wound to naturally resume re-epithelialisation but without an ‘overwhelming’ provision of autologous cells. This may also provide biological clues as to why skin grafts do not always demonstrate optimal tissue “take” and efficacy.

Collectively, our *in vivo* findings confirm that treatment with VeritaCell ACS at a 1:20 donor-to-wound ratio (and even more so a 1:10 ratio) not only shows significant acceleration of skin neo-epithelisation, but also enhances the quality of wound recovery by demonstrating a reduction in scarring-associated tissue architecture. Traditionally, however, it has been suggested that although excisional wound models in rats provide valuable insights into skin repair (23), they differ from human healing processes in key anatomical and physiological aspects. Rodent wounds have been postulated to close mainly *via* contraction and less so by re-epithelialization, with the subcutaneous *panniculus carnosus* muscle driving rapid contraction, a feature absent in humans. Thus, it has often been suggested that ‘wound splinting’ is required (24) to permit assessment of scarring-associated responses. More systematic studies, however, have provided evidence that such excisional wounds provide a valid and reproducible wound model (21) and this permits observations on wound healing quality to be made. Attesting to the ability of such models to provide clues on scarring-associated responses, is evidence generated by several *in vivo* rat studies demonstrating how a variety of wound response-modulating treatments influence cutaneous thickening (25-28). Our observations on the ability of ACS suspensions to suppress excessive, scarring-associated neo-epithelial layer thickening provides further support for the suitability of these *in vivo* models.

Finally, our study highlights an important advantage of the ACS methodology, which is the requirement of a much smaller amount of donor skin to enhance wound healing on a larger skin area. Unlike using STSG, which requires donor-to-wound site ratios of 1:1 (or at best 1:2), ACS application clearly provides wound benefit by requiring remarkably less donor skin. Of note, it has previously been suggested that even a 1:80 donor site-to-wound site ratio might be clinically beneficial (8). We would nevertheless exercise caution in ‘compromising’ the benefit of the ACS approach by ratios that are higher than 1:20. Not only are our *in vivo* observations supportive of a potentially ‘optimal’ donor-to-wound site ratio (and different such ACS application ratios may be utilised by clinical partitioners depending on the severity of the wounds to be treated), but also they may provide an explanation as to why, so far, ACS-based therapy has not always achieved the desired clinical promise (also in the context of method cost *versus* clinical benefit) (18).

In conclusion, following on from our previous published findings on the beneficial biological properties of ACS-associated cell populations (provision of viable cell populations and a wound healing-enhancing secretome), we now demonstrate clear wound healing benefits, as VeritaCell suspension-treated skin exhibited accelerated healing rates (even in the context of an acute healing response), as well as demonstrating enhanced ‘quality’ of wound healing by minimising scarring-associated tissue thickening. These preclinical observations strongly support the clinical use of VeritaCell ACS for enhancement of wound responses in critical wounds requiring rapid closure acceleration (such as burns), and also offer insights on the promise of the ACS approach in providing a medical solution that will facilitate more aesthetically favourable healing outcomes.

## Author contributions

MP: study conceptualisation, experimental planning, conducted experiments, data analyses and interpretation, co-led preparation and finalisation of manuscript. OV & JU: conducted experiments. VP: data analyses and interpretation. AS: study conceptualisation, experimental planning, funding acquisition. BJ: experimental planning, conducted experiments, data interpretation, co-led finalisation of manuscript. NTG: study lead, study conceptualisation, experimental planning, data interpretation, funding acquisition, led preparation and finalisation of manuscript.

## Acknowledgments

The authors are indebted to Ms Elga Poppela for her unrivalled technical support with animal care during the *in vivo* studies. We also thank Dr Kristine Nevidovska and her team (Academic Histology Laboratory Ltd, Riga, Latvia) for performing the histology work.

